# State-Dependent Stimulation for Seizure Termination in a Morris–Lecar Network: A Reinforcement Learning Approach

**DOI:** 10.64898/2026.11.11.765955

**Authors:** Veronika Házi, Marcell Stippinger, Tamás Kiss, Zoltán Somogyvári

## Abstract

The efficacy of neurostimulation for seizure termination depends not only on the stimulus itself but also on the dynamical state of the epileptic network. Here, we develop a dynamical-systems-informed reinforcement-learning framework that learns when to deliver a predefined pulse to terminate seizure-like activity. We simulate a sparsely connected network of 100 extended three-dimensional Morris–Lecar neurons. Bifurcation analysis of the single-neuron model identifies bistable regimes and provides a mechanistic basis for considering transitions between resting and oscillatory activity. The learning agent uses semi-gradient Q-learning with tile coding and receives a two-dimensional time-delay embedding of either the population-averaged membrane potential or a derivative-based proxy for extracellular potential. It learns a state-dependent policy that selects between stimulation and no stimulation, without modifying the pulse waveform. Across 30 independently generated test networks, both learned policies markedly reduced median seizure duration compared with trigger-only and state-independent stimulation controls. The two learned policies performed comparably, indicating that the estimated extracellular potential proxy retained sufficient dynamical information for control in this model. These results provide a proof of concept that combining dynamical-systems-based state reconstruction with online reinforcement learning can identify effective stimulation timing for terminating seizure-like activity.

## 1 Introduction

Epilepsy is a major clinical burden, with recent studies estimating approximately 50 million epileptic patients worldwide (GBD Epilepsy Collaborators 2025; Zhao et al. 2026). Despite continuous research in antiseizure drug development, one third of these patients remain drug-resistant and need alternative therapeutic approaches (Grávalos et al. 2025).

Neurostimulation is a promising tool for treating drug-resistant epilepsy (Berényi et al. 2012; Barcsai et al. 2026). Although alternative modalities, such as focused ultrasound (Chen et al. 2020), are being investigated, electrical stimulation remains the most established approach in clinical practice (Rao 2025). Despite its common application, finding the right stimulation parameters remains a major challenge for effective treatment. While finding the correct stimulation waveform is a challenge in itself, we must consider that its effect may also depend on the state of the nervous system. This raises the question of whether a short, simple pulse could be sufficient to terminate a seizure if delivered at the right time.

Experimental results (Osorio and Frei 2009; Barcsai et al. 2026) as well as model simulations (Suffczynski et al. 2004; Taylor et al. 2014) suggest that a single pulse could indeed be enough to stop a seizure under certain circumstances. These studies also showed that the efficacy of stimulation depends on the instantaneous phase of the epileptic oscillation, as observed in EEG. Stimulation timing is also emphasized in recent reviews of responsive neurostimulation (RNS), a closed-loop electrical technique that is already used in medical practice (Rao 2025). These results highlight the importance of the phase of the epileptic oscillatory system when applying stimulation. While the phase is undoubtedly an important state variable, it is typically insufficient on its own to fully describe the state of the system, making the outcome of stimulation potentially ambiguous. Thus, a more complete description of the internal state is necessary to implement a state-dependent stimulation protocol. Once a sufficient state representation has been identified, an optimal stimulation protocol could be defined as an optimal policy over these states.

Izhikevich (2007) described the dynamics of excitable systems operating near a bifurcation point, where the resting state, represented by a stable fixed-point attractor, loses stability and gives way to a more dynamic, periodic (or chaotic) attractor corresponding to the excited state. This perspective provides a framework for classifying seizure onsets and offsets according to the bifurcations involved.

Jirsa et al. (2014) and Saggio et al. (2020) classified seizure dynamics into different types, termed dynamotypes, based on the bifurcation types involved and the geometrical relationship between the fixed point and the periodic attractor in state space. Using this dynamotype classification, Szuromi et al. (2023) investigated the effects of short biphasic stimulation pulses on two-dimensional dynamical systems belonging to different classes. They concluded that the stimulation phase and amplitude required for effective seizure termination depend on the dynamotype of the system. Since finding the optimal stimulation parameters is non-trivial even in these relatively simple systems, determining them analytically in more complex, higher-dimensional systems is even more challenging.

Here, we investigate whether reinforcement learning (RL) can learn an effective seizuretermination policy through interaction with a network exhibiting seizure-like activity.

Reinforcement learning is a machine learning approach in which an agent learns from experience rather than from labeled data. The agent can interact with an environment, observe the results of its actions, and receive rewards that indicate how desirable the actions were. This framework is well suited to the challenge of developing a state-dependent, adaptive stimulation strategy. In this context, the epileptic brain could serve as the environment, whose states should be at least partially observable to the agent, while the reward is determined by the rapid termination of the seizure with minimal intervention.

Several studies have already applied RL to similar problems. In Parkinson’s disease research this is a well-explored area with a large number of recent publications (Lu et al. 2020; Cho et al. 2024; Zhao et al. 2025). However, in epilepsy research this field remains limited. In a series of publications, Pineau et al. (Guez et al. 2008; Pineau et al. 2009; Panuccio et al. 2013) demonstrated policy evaluation by offline training of an RL agent on previously recorded data in an *in vitro* epilepsy model. Although the learnt policy did not result in shorter seizures than periodic stimulation, it achieved a similar result with less intervention. A few papers have implemented online RL algorithms to control simulated epileptic models: Nagaraj et al. (2017) implemented an online algorithm using the Epileptor model (Jirsa et al. 2014). More recently, the cortical-thalamic neural-mass loop model was also assessed under different RL systems, in which online learning and active feedback were feasible (Dan et al. 2026). However, in these simulated systems, seizures were suppressed only during the periods when continuous control was turned on, whereas both experimental results and model simulations suggest that a single well-timed pulse could be enough to terminate a seizure (Osorio and Frei 2009; Suffczynski et al. 2004; Szuromi et al. 2023). Consequently, RL studies have so far failed to demonstrate that RL can learn a stimulation policy that terminates the seizure by returning the system to the resting state.

The aim of our work is to develop an *online* RL system capable of learning an effective stimulation policy for terminating epileptic seizures. We aim to demonstrate the potential of this approach in a simulated environment while maintaining compatibility with real EEG recordings for future applications. For this purpose, we create a model of a potentially bistable epileptic network and test whether the stimulation policy can drive the system back into the normal activity regime with a few properly timed stimulation pulses, or even a single one. We target a fundamentally different time scale than offline RL works (Guez et al. 2008; Pineau et al. 2009; Panuccio et al. 2013): we explicitly characterize the instantaneous state of the epileptic system and determine the appropriate timing of individual stimulation pulses on the millisecond scale. This approach takes into account and exploits the oscillatory dynamics of the network, providing the potential to terminate seizures more rapidly with less intervention.

### 1.1 Modeling Considerations

The neural network model we developed (Barcsai et al. 2026) aims to capture the dynamical changes that lead to seizure onset or enable the system to escape from a seizure state, without attempting to reproduce all the details of the underlying neurophysiology.

Jirsa et al. (2014) classified seizure dynamics according to the bifurcation types associated with seizure onset and offset. Assuming that simultaneous transitions across critical points in multiple parameters are unlikely, four possible codimension-one bifurcation types can describe the transition from a fixed-point attractor to a periodic attractor (Izhikevich 2007), and four types describe the reverse transition. Together, these yield 16 possible onset-offset bifurcation pairs. Furthermore, Saggio et al. (2020) and Szuromi et al. (2023) extended this classification and showed that the geometrical relationship between the periodic limit-cycle attractor and the fixed point in state space is a key factor in seizure termination. Considering two-dimensional systems, two major classes have been identified: systems in which the fixed point lies inside the limit cycle, referred to as LC-big (LCb) systems, and systems in which the fixed point lies out-side the limit cycle, referred to as LC-small (LCs) systems. They showed that seizures are easier to terminate in LCs systems than in LCb systems: a sufficiently large pulse can terminate seizures in LCs systems with high probability, whereas termination with a smaller pulse requires precisely tuned stimulation within a specific phase range.

Note that, although this classification works well for two-dimensional systems, the geometrical relationships between limit cycles and fixed points become more complex in higher-dimensional systems. In three or more dimensions, a fixed point cannot be classified as lying inside or outside a limit cycle in the same way as in two dimensions. Instead, the relative geometry of the basins of attraction of the two attractors becomes the relevant factor.

Although simplified neural mass models can describe epileptic phenomena using only a handful of differential equations, they cannot account for network-level properties of epileptic seizures, such as neuronal synchronization and desynchronization. Since stimulation can significantly affect network synchrony, we constructed a network of sparsely interconnected simple neurons. This approach allows us to analyze the dynamics of individual neurons in detail while the RL agent observes and interacts with the population-level activity.

## 2 Methods

### 2.1 Single-Cell Model

Our epilepsy simulations were based on a network of modified Morris–Lecar neurons (Morris and Lecar 1981). The network consisted of one hundred sparsely connected three-dimensional neurons governed by the following system of differential equations:

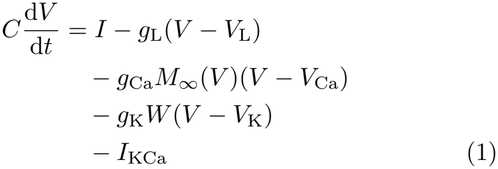

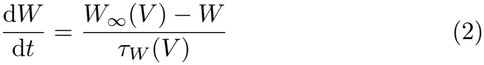

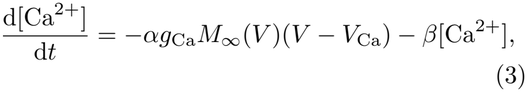

Where

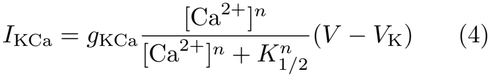

and the steady-state and time-constant functions were:

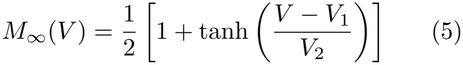

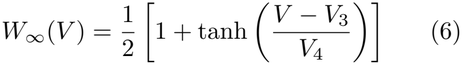

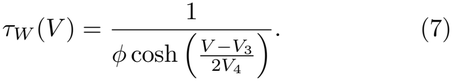

The original planar system was extended with slow [Ca^2+^] dynamics to produce seizure-relevant bursting (Ermentrout and Terman 2010; Azizi and Mugabi 2020). The model parameters are presented in Table A1.

#### 2.1.1 Bifurcation Analysis of the Single-Cell Model

The dynamics of the single-neuron model were examined in the time domain and in phase space under different conditions. To characterize the dynamical behaviour of the model, we performed bifurcation analysis using the XPPAUT software package (Ermentrout 2002). We used the injected current as the bifurcation parameter, as it can drive the behaviour of individual neurons. The equilibrium points were found by a Monte Carlo search. We determined their stability by the eigenvalues of the Jacobian matrices. Equilibrium branches were subsequently traced using numerical continuation through the integrated AUTO module, followed by the continuation of periodic solutions.

In some regions, the software struggled to consistently classify bifurcations, especially at the termination of unstable periodic branches. We inspected these points in more detail using the change in the oscillation period and the stability of nearby singular points. If the oscillations approached a hyperbolic saddle and their period diverged logarithmically as the distance from the bifurcation point shrank, we declared a saddle-homoclinic (HC) bifurcation. For a saddle-node on invariant circle (SNIC) bifurcation, an eigenvalue with zero real part should exist and the oscillation period should scale as (Izhikevich 2007):

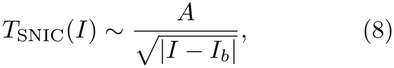

while for a saddle-homoclinic bifurcation the corresponding relationship is:

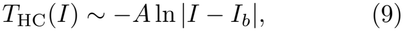

where *I*_*b*_ corresponds to the current at the bifur-cation point. The most likely bifurcation type was identified by combining the eigenvalue-based classification of nearby equilibria with the goodness of fit of the theoretical period-scaling relationships.

This dynamical analysis was performed on a single neuron only, whereas seizures were evoked by the complex interactions between multiple neurons. Hence, the results of this analysis are not conclusive for the network, but they can still provide meaningful insights into the potential mechanisms underlying seizure generation and termination.

### 2.2 Network Model

The final network consisted of one hundred extended Morris–Lecar models with sparse interconnections. The model was integrated using the forward Euler method. Besides the synaptic input from other cells, all neurons had some baseline activity. Therefore the input current *I* comprised two components. For the *j*th neuron:

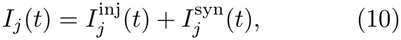

where *I*^inj^ and *I*^syn^ represent the background excitation and the synaptic input, respectively. The synaptic input represents the interaction between the neurons and is defined as:

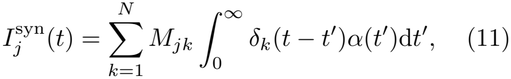

where *M*_*jk*_ denotes the *jk*^th^ element of the connection matrix *M* , and *δ*_*k*_(*t*) is the Dirac delta centred on the spike times of neuron *k*. In the simulations, time was discretized with the step d*t*, a spike was registered as a binary event when *V* crossed 0 mV from below, and the integral was evaluated as a sum of the values of the alpha function. Alpha functions represented synaptic currents, which were generated by each node upon activation and were added to all connected neighbours:

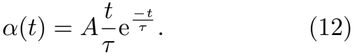

For each simulation, the binary connection matrix *M* was generated by independently setting each element to 1 with probability *p* = 0.1. Thus, the expected number of connections was 1000 in the 100 × 100 connection matrix.

*I*^inj^ represents the background excitation:

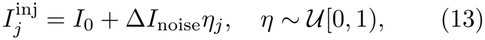

where *I*_0_ was the baseline injected current, Δ*I*_noise_ controlled the magnitude of the static offset per neuron, and *η*_*j*_ was a random variable drawn from a uniform distribution independently for each node.

Seizure-like activity was induced by stimulating a single neuron at time *T*_init_. To characterize the resulting collective dynamics of the network, we calculated the population-averaged state variables, defined as follows:

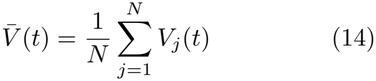

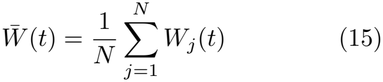

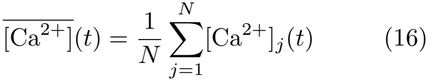

The membrane potential was used to calculate the Golomb–Rinzel synchrony index in the network as defined in Golomb (2007). The variances were only measured during seizure-like activity, from the first time 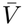 exceeded 0 mV after the initial stimulation until it had remained below 0 mV for 1 s. As a control, the same index was computed after each neuron’s trace was circularly shifted by an independent random amount. For independent neurons the index is expected to be about 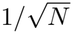 , where *N* = 100 is the number of neurons in the network. Values are reported as mean SD over *n* = 30 networks.

Table A2 summarizes the parameters of the network. It also includes the time step and the total simulation time.

### 2.3 Definition of Synthetic EEG

The average membrane potential 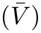 across all neurons was used to define a proxy for the extracellular potential, an approximation of an experimentally accessible variable:

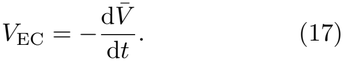

Note that, due to the dipolar nature of the extra-cellular potential, the sign of this approximation depends on the geometrical arrangement of the neurons and the recording electrode system.

### 2.4 Definition of State Representation

Successful reinforcement learning requires a proper state representation as input. Because the internal state variables of the network 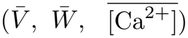 cannot be measured directly in a realistic environment, we used Takens’ time-delay embedding theorem (Takens 1981) to reconstruct the state space. The theorem states that the attractor of a dynamical system can be reconstructed, up to topological equivalence, from the time series of any smooth scalar-valued observation function of the state variables. Topological equivalence means that the neighbourhood relations between states are preserved: the reconstructed attractor is a transformed version of the real one, but similar states remain close to each other and the overall topological structure is maintained. In other words, we can use a quantity directly accessible from measurements, such as an EEG time series (approximated here as *V*_EC_), take its delayed versions ((*V*_EC_(*t*), *V*_EC_(*t* − *τ*), *V*_EC_(*t* − 2*τ*), …)), and reconstruct a state space without losing major dynamical features. Accordingly, *V*_EC_ was used to construct one of the state representations. Since the membrane potential (*V*) was directly accessible from the model and exhibited a less noisy state-space structure, we used it as a reference state representation.

Although the dimension of the system is higher than two, using only two state variables was sufficient for learning and has multiple advantages for applications. First, it was apparent that the main dynamical components of the trajectories, such as the limit cycle and the resting point, remained distinguishable even after the 2D projection. A lower-dimensional state representation can also reduce the complexity of the learning problem and potentially improve sample efficiency. It could also be beneficial in real-time systems, where a single time-delayed coordinate introduces only a short latency and can be calculated efficiently.

### 2.5 Reinforcement-Learning Algorithm

The Reinforcement-Learning (RL) framework is based on agent-environment interactions. In our work, the Morris–Lecar network serves as the environment in which the agent learns. We implemented a simple agent using a semi-gradient Q-learning algorithm with tile-coding and *ϵ*-greedy action selection (Sutton and Barto 2018). It observes the time-delay embedded states (*S*) of the network and interacts with it through a discrete action space consisting of two actions, *A*_0_ (no stimulation) and *A*_1_ (stimulation). The stimulation waveform is fixed in advance. The agent only decides when to deliver it. The proposed frame-work with the predefined stimulation pattern is displayed in Figure 1.

**Fig 1:**
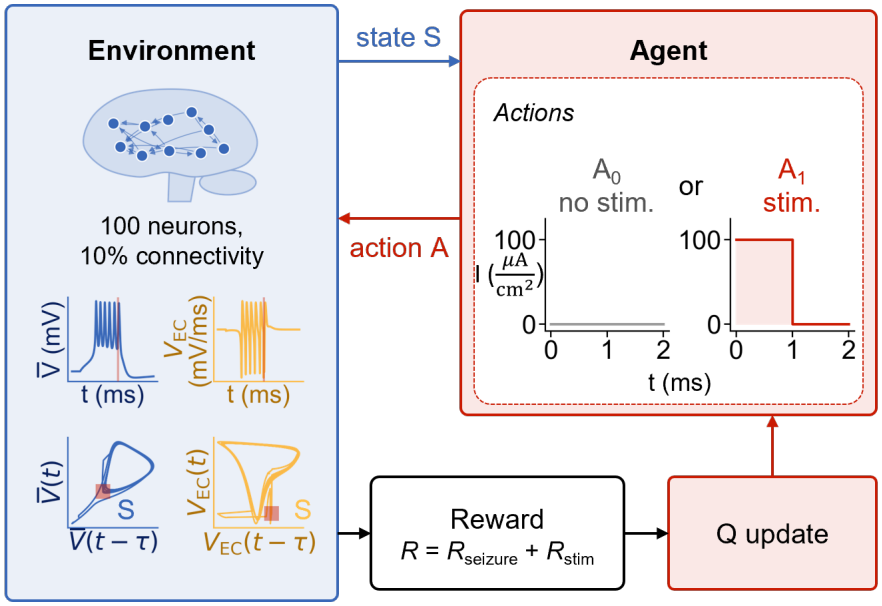
Framework of the reinforcement learning algorithm for seizure termination in a simulated neuronal network. The agent observes the state *S* of the network, represented by a 2D time-delay embedding of either the mean membrane potential 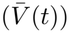 or the extracellular potential proxy (*V*_EC_). It also chooses whether to stimulate (*A*_1_) or not (*A*_0_). The stimulation policy is then updated based on the received reward.

Tile coding helps the agent to be sample efficient and to handle large, continuous state spaces. The space is divided into discrete units, and this division is shifted multiple times to produce further discretizations, called tilings. The units of the tilings are the tiles. Instead of storing action-values (*Q*(*S*_*t*_, *A*_*t*_)) for the infinitely many states, action-values are assigned to individual tiles. When the agent makes an observation, it first identifies all tiles that contain the observed state. These are the active tiles. The action-value of the state is defined as the sum of the weights of the active tiles for the corresponding action:

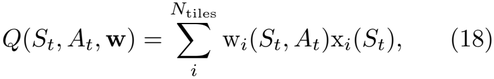

where *N*_tiles_ is the number of tiles, w_*i*_ is the weight of tile *i* and x_*i*_(*S*_*t*_) is the *i*^th^ element of the feature vector of state *S*_*t*_, defined as:

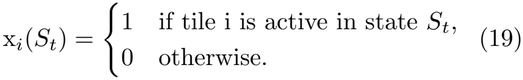

The semi-gradient Q-learning algorithm updates the tile weights at every iteration using the following update rule:

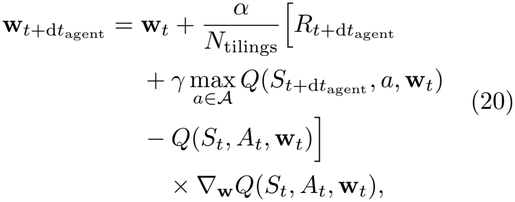

where *α* is the learning rate (also called step size) and *γ* is the discount factor. For linear function approximation, the gradient simplifies to the feature vector:

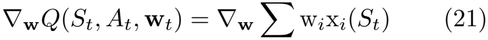

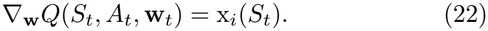

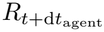 is the reward received for action *A*_*t*_. We calculated the reward from the average network activity 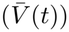 as the time elapsed since the last seizure, scaled by the maximum seizure duration. The time of seizure cessation was defined as the last time at which the average membrane potential was positive:

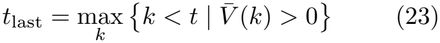

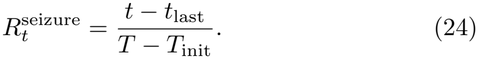

An additional penalty was applied whenever the stimulation action was performed. Its size was chosen arbitrarily, but it was scaled in the same way as the seizure duration to keep the reward bounded:

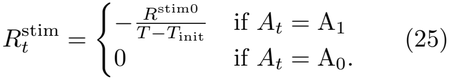

The total reward is thus:

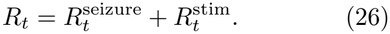

After every interaction, the agent updates the tile weights and selects the next action. Actions were selected with an *ϵ*-greedy policy: with probability 1 − *ϵ* the agent selected the action with the highest action-value, and with probability *ϵ* it explored by choosing an action at random. To encourage exploration at the beginning of training and exploitation at the end, the parameter *ϵ* was reduced continuously according to the following exponential decay:

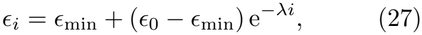

where *i* is the index of the training network and *ϵ* is kept constant during the episodes on the same network (see Section 2.5.1). Parameters were chosen such that exploration and exploitation were initially equally likely, while by the end of training the agent practically always chose exploitation.

#### 2.5.1 Training and Evaluation

We conducted two rounds of training and evaluation. Initially, we used 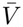 for the state representation and later repeated the experiment using *V*_EC_. The training and testing procedure was the same in both conditions and is described in detail below.

The agent was trained over 50 sessions, each consisting of 70 episodes. A session was defined by providing a new network to the agent, while episodes were repeated runs on the same network with different a gent a ctivation t imes. A session started by generating a new connectivity matrix and fresh baseline input currents. The *ϵ* parameter was also set at the beginning of each session using Eq. 27. Each episode started by triggering the network at time *T*_init_ to evoke a seizure, after which the agent was activated with a delay (*t*_delay_). After activation, the agent interacted with the network in discrete steps. At every time step it chose an action, carried it out in the environment, and received a new state and the corresponding reward. This information was used to update its weights and to select the next action. This cycle continued until the simulation time of the network was over.

The delay of agent activation comprised the duration of the initial stimulation (*t*_init_), a fixed delay allowing the stimulation to take effect (*t*_eff_), and a noise term (*δ*). The noise was drawn from a uniform distribution between 0 and *T*_jit_.

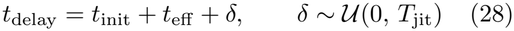

To analyse learning performance, we collected the accumulated rewards per episode (returns), the number of stimulations and the duration of seizures. The mean and standard deviation (SD) of these metrics were computed across episodes.

After training, the learnt weights of the agent were fixed and its policy was switched to greedy action selection, that is, it always selected the action with the higher action-value. Testing was performed over 30 new sessions with 10 episodes each. Returns, stimulation numbers and seizure durations were collected again. Seizure durations were compared across four conditions. First, the original network was simulated without agent activation. Second, a blind agent performed state-independent stimulation: it was assigned the median number of stimulations and the median seizure duration obtained from the trained agent, and it distributed the corresponding number of stimulation pulses randomly over this time interval, independently of the network state. Lastly, the network was evaluated using the policies learned on the two state representations, 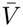 and *V*_EC_.

For each test network, the seizure duration of a condition was summarised by the median over its ten episodes, giving one value per network (*n* = 30 per group). Conditions were compared with two-sided Mann–Whitney U tests on every pair, with Holm–Bonferroni correction over the six comparisons. Non-parametric tests were used because seizure durations are bounded and right-skewed.

The parameters of the reinforcement-learning algorithm are listed in Table A3.

### 2.6 Use of AI

The Morris–Lecar network model and the reinforcement-learning algorithm were designed and implemented by the authors without AI assistance. Claude (Anthropic) was used to debug the code, write plotting code that draws the figures from the simulation data, and implement the statistical tests and the synchrony analysis according to methods specified by the authors. All AI-assisted code, analyses and text were checked and revised by the authors, who take full responsibility for the content.

## 3 Results

### 3.1 Bifurcation Analysis of Single-Cell Model

The single-cell model exhibited different behaviours under different input currents. To investigate the underlying mechanism of these transitions, we examined its dynamical properties with XPPAUT. With *I* = 0 *μ*A*/*cm^2^ the model only had one equilibrium point at (*V* = −59.53 mV, *W* = 2.21 *×* 10^−4^, [Ca^2+^] = −0.219) with eigenvalues of *λ*_1_ = −0.095, *λ*_2_ = −1.007, *λ*_3_ = −0.005. Because all eigenvalues are negative and real, the equilibrium is classified as a stable node. This corresponds to the resting state, where the neuron would remain in the absence of external stimuli. In the network, all neurons received at least the baseline current *I* = *I*_0_. Hence, the model was also characterized around that value. Table B4 shows the three fixed points that emerged at *I* = *I*_0_ = 41*μ*A*/*cm^2^. Based on the eigenvalues, the first point (#1) is a saddle point, the second (#2) is a stable node, while the last one (#3) is a saddle-focus, highlighting the diversity of local dynamics in this region. Lastly, we inspected the neuron at *I* = *I*_init_ = 45*μ*A*/*cm^2^, the current used to trigger seizure-like activity in the network. The software found only one fixed point (*V* = 3.81 mV, *W* = 0.28, [Ca^2+^] = 59.09) with eigen-values of *λ*_1,2_ = 2.4 10^−4^ *± i*0.379, *λ*_3_ = 0.005. This classifies it as a saddle-focus, indicating that nearby trajectories tend to spiral away from this point.

The variation in the number and stability of the fixed points with the input current suggested the presence of bifurcations. To confirm this intuition, we performed a bifurcation analysis. To illustrate how the analysis translates into behaviour, we present the response of the single-cell model to an applied current of *I* = *I*_init_ = 45*μ*A*/*cm^2^. Fig.2(a–h) shows the resulting time series and trajectories, while Fig.2(i,j) depicts the attractor branches obtained from the bifurcation analysis.

In the simulation, the neuron quickly converged to a stable periodic oscillation, as predicted by the monostable limit-cycle attractor in the bifurcation diagram of Fig. 2(j). The structure of the cyclic attractor is visible in the phase plane and in the embedded state spaces in Fig. 2(f–h). Although no stable resting state exists at this current level, the stable equilibrium at *I* = *I*_0_ = 41 *μ*A*/*cm^2^ is marked for reference to indicate where the resting state is located under baseline conditions.

**Fig 2:**
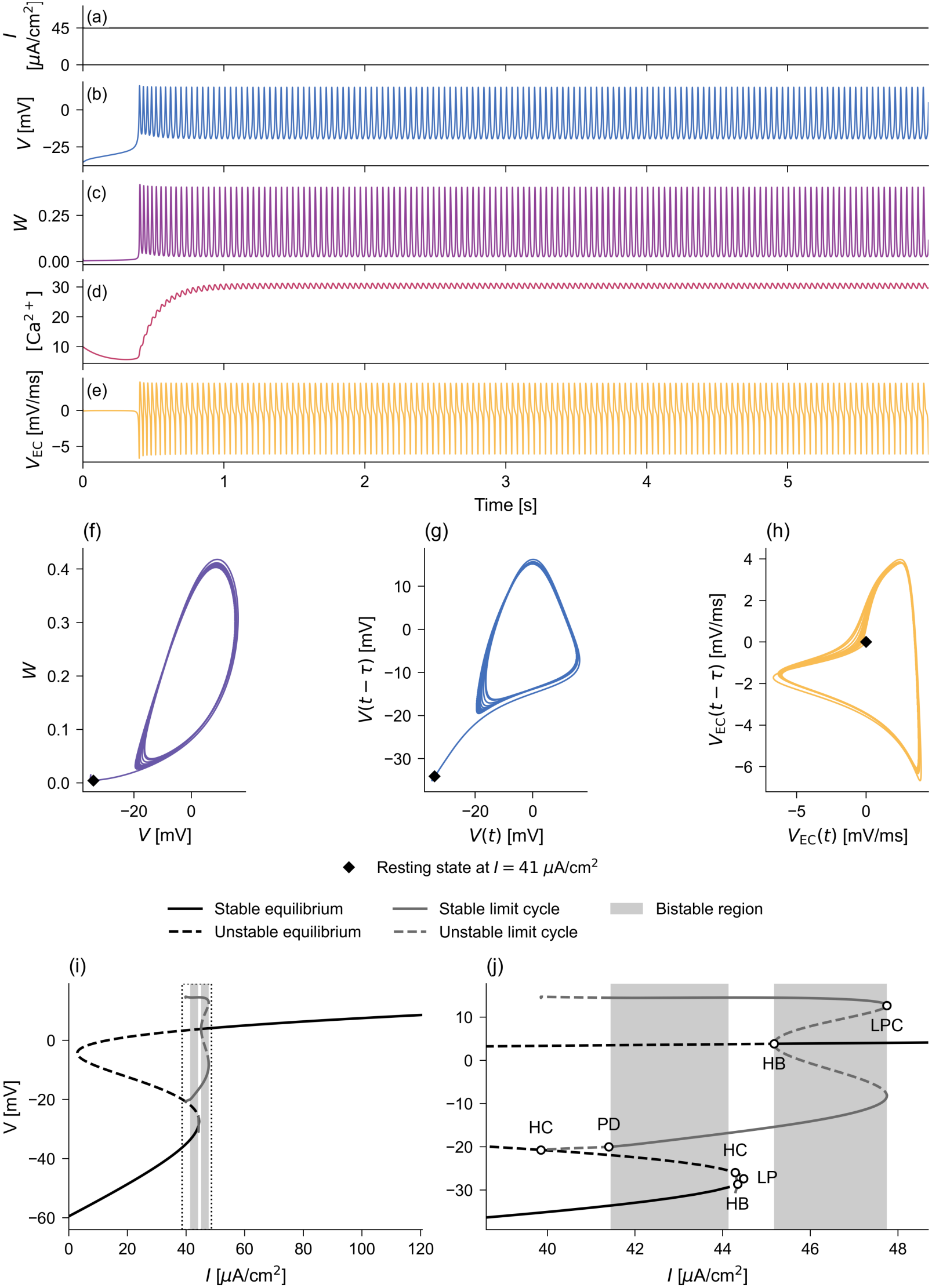
Dynamics and bifurcation analysis of the 3D Morris–Lecar neuron. (a–e) Time courses of the input current (*I* = 45 *μ*A*/*cm^2^) and model variables (*V* , *W* , [Ca^2+^], *V*_EC_). (f–h) Trajectories in the *V* –*W* phase plane and delay-embedded state spaces, with the resting state at *I* = 41 *μ*A*/*cm^2^ marked by black diamonds. (i–j) Bifurcation diagrams showing the full parameter range and an enlarged view of the bifurcation points, respectively. Solid and dashed lines indicate stable and unstable branches, while black and gray represent equilibria and limit cycles, respectively. Shaded areas highlight bistable regions. Bifurcations: fold of limit cycles (LPC), saddle-node (LP), Hopf (HB), period-doubling (PD), and saddle-homoclinic (HC).

Bifurcation analysis revealed bistable regions at 41.4*μ*A*/*cm^2^ ≤ *I* ≤ 44.34*μ*A*/*cm^2^ and 45.17*μ*A*/*cm^2^ ≤ *I* ≤ 47.75*μ*A*/*cm^2^, highlighted in Figure 2(i,j). This confirmed that, with the selected baseline current parameters (see Table A2), most neurons in the network started in a bistable regime, 41*μ*A*/*cm^2^ ≤ *I* ≤ 44*μ*A*/*cm^2^. Around the bistable regions, several different types of bifurcation points were identified by XPPAUT. They are marked and labelled in Fig. 2(j).

All bifurcations were classified by XPPAUT except the termination points of the unstable oscillatory branches. These points were crucial for understanding how the neuron can lose its periodic behaviour, i.e. how a seizure can terminate. The period increased greatly around both points, suggesting either a saddle-node on invariant circle (SNIC) or a saddle-homoclinic (HC) bifurcation. The traced singular points were consistently hyperbolic saddles with two negative and one positive eigenvalue (see Table B5 and Table B6), which favoured a HC rather than a SNIC bifurcation. We also fitted the two characteristic period-scaling functions to the period change (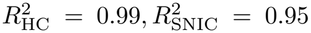 for *I* = 39.85 and 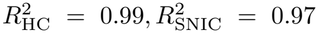 for *I* = 44.29, see Figures B1 and B2), which favoured a HC point in both regions.

Overall, the dynamics of the modified Morris– Lecar neuron can be characterized by a sequence of transitions as the input current increases. At low input currents, the system is monostable, exhibiting a single stable equilibrium point. As the input increases, the system enters a bistable regime in which a stable equilibrium coexists with a stable limit cycle. With further increase in the input current, the equilibrium loses stability, leaving the periodic limit cycle as the only stable attractor. At even higher input currents, a new stable equilibrium emerges above 0 mV, initially coexisting with the limit cycle before becoming the sole stable attractor.

### 3.2 The Morris–Lecar Network exhibits seizure dynamics

The Morris–Lecar network exhibited seizure-like episodes when triggered. To induce seizures in a controlled manner, an initial current of amplitude *I*_init_ = 45*μ*A*/*cm^2^ and duration *t*_init_ = 0.1 s was applied at time *T*_init_ = 1 s to a randomly selected neuron in every network. This usually resulted in seizure-like activity at the network level.

We demonstrate the typical seizure pattern of our model in Figure 3. This network is an example from the training set and was arbitrarily chosen to illustrate some important concepts. The time courses of the averaged state variables in Fig. 3 (a–c) reveal how the network produces seizure-like oscillations after the initial stimulus and how these can terminate spontaneously later. The approximated extracellular potential (*V*_EC_) is also depicted (Fig. 3 d), as it represents what could be measured in an experimental setting. Fig. 3 (e) portrays the seizure dynamics in a phase plane constructed from two of the original variables (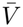 and 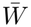). During a seizure, the system follows a trajectory along the tilted elliptical structure until it returns to the resting point, located in the lower left corner of the phase plane. The time-delay reconstruction from the membrane potential reveals a matching structure (Fig. 3 f). The attractor reconstructed from *V*_EC_ (Fig. 3 g) shows a similar picture, but the structure is less clear and noisier.

**Fig 3:**
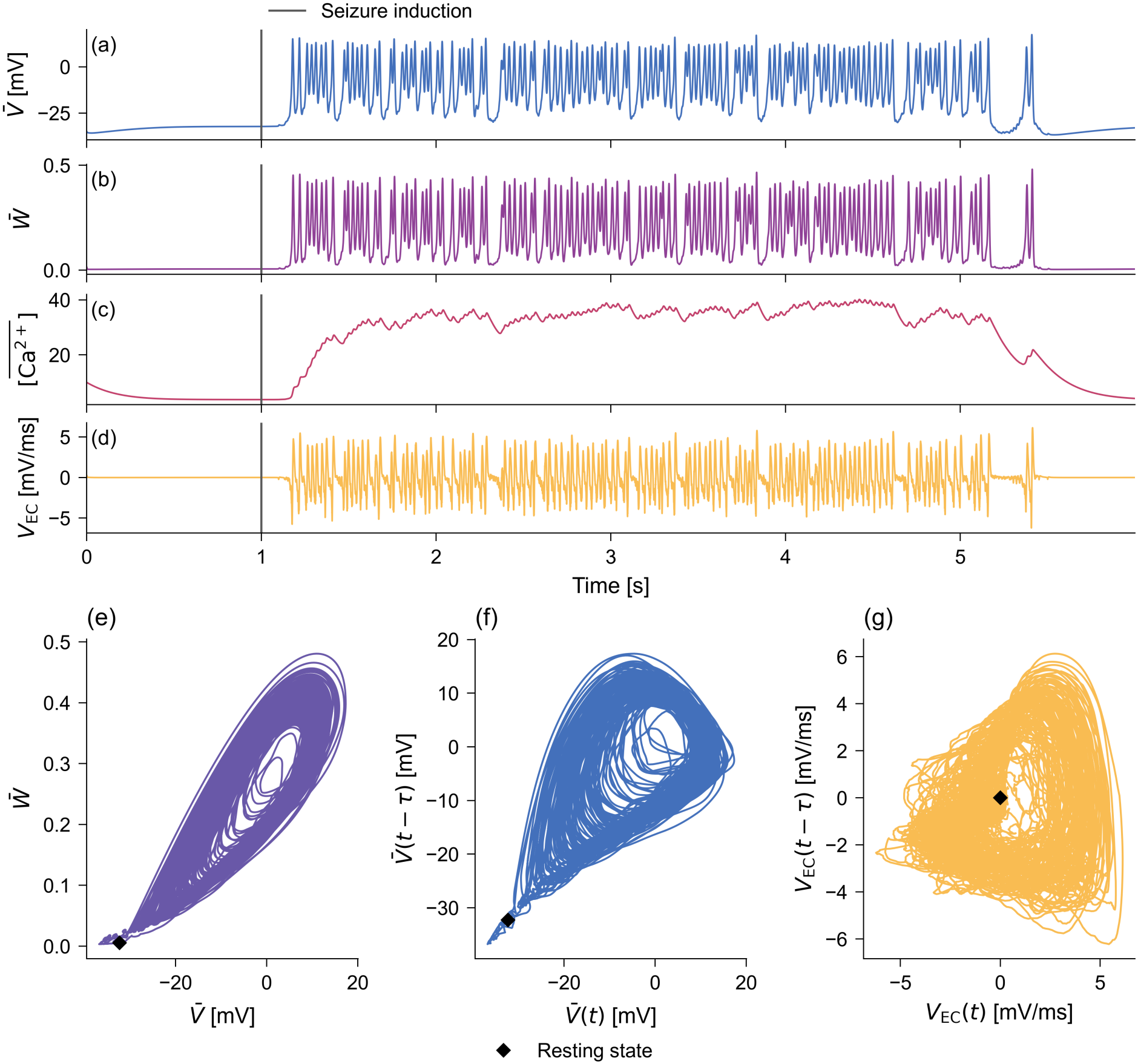
An example episode from the training set showing the activity of the Morris–Lecar network: seizure-like behaviour when triggered, followed by spontaneous termination. The signals are averaged across neurons (*N* = 100) to represent the whole network. (a) Time course of the mean membrane potential 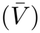. (b) Time course of the mean recovery variable 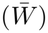. (c) Time course of the mean intracellular calcium concentration 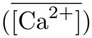. (d) Time course of the extracellular potential proxy (*V*_EC_). (e) Network trajectory in the 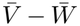 phase plane. (f) Network trajectory in the delay-embedded state space constructed from the 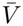 membrane potential. (g) Network trajectory in the delay-embedded state space constructed from the *V*_EC_ extracellular potential proxy. The resting state is marked with a black diamond and is calculated from the network means averaged over the 0.8–1 s period.

Although the bifurcation analysis was performed on an isolated neuron, the network exhibited qualitatively similar dynamics. Barcsai et al. (2026) estimated the intrinsic dimension of this attractor using a *k* -nearest-neighbour estimator (Benko et al. 2022). The estimated median dimension of the population activity was 3.07 ± 0.38 (median absolute deviation). This means that the large number of potential degrees of freedom of the whole network were trapped, through population synchrony, in a low-dimensional oscillation. The cyclic attractor retained a similar structure in both the phase plane and the embedded state spaces, while the resting state occupied a similar position. The latter was expected, as rest in the network corresponds to little or no synaptic input to most neurons, in which case the initial conditions place the neurons in the basin of attraction of the fixed point (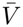 ≈ −32 mV). At *T*_init_, an increased current (*I*_init_) was applied to a single neuron, which placed it in the regime of the monostable periodic attractor, as seen in Fig. 2. The resulting spiking activity could propagate through the synaptic connections, increasing the input currents of other neurons and potentially driving them into the bistable regime or across the Hopf bifurcation threshold. As more neurons are recruited into sustained firing, recurrent synaptic excitation may give rise to prolonged network-wide oscillations resembling the seizure-like activity observed in epilepsy. During a seizure, neurons tend to synchronise, which is one of the most characteristic features of epilepsy. The Golomb–Rinzel synchrony index of the membrane potentials during seizure-like activity, measured across *n* = 30 networks, was 0.730 ± 0.033. For the randomly shifted traces the index fell to 0.100 ± 0.005. This indicates strong, but not absolute, synchrony during these episodes. These observations supported the suitability of the Morris–Lecar model for investigating seizure-like activity at the network level.

### 3.3 The RL agent learns to terminate seizure-like activity through state-dependent stimulation

The agent was trained over 50 sessions with 70 episodes each. Performance was measured by the accumulated rewards (returns), the number of stimuli delivered, and the duration of the seizures across sessions. The learning curves, depicted in Figure 4 (a–c), demonstrate convergence in all three metrics. The returns tended to increase (Fig. 4 a), while the number of stimulations and the seizure lengths decreased continuously (Fig. 4 b,c). These observations indicate that the agent progressively learned to terminate seizures more rapidly and with fewer interventions, collecting increasing rewards. The number of stimulations converged earlier than the seizure lengths, suggesting that the agent learned the correct timing towards the end of training. Note that no minimum number of stimulations was imposed. The agent could decide not to intervene at all, which was the correct choice if there was no seizure or if it ended spontaneously after a short period. In contrast, seizure duration had a lower bound due to the delay between seizure initiation and agent activation. Consequently, seizure duration could not converge to 0 s even if the agent terminated the seizure immediately upon activation. Since the activation delay varied across episodes, this lower bound was not constant. To estimate the value to which the agent could converge, we calculated the average activation delay across episodes.

**Fig 4:**
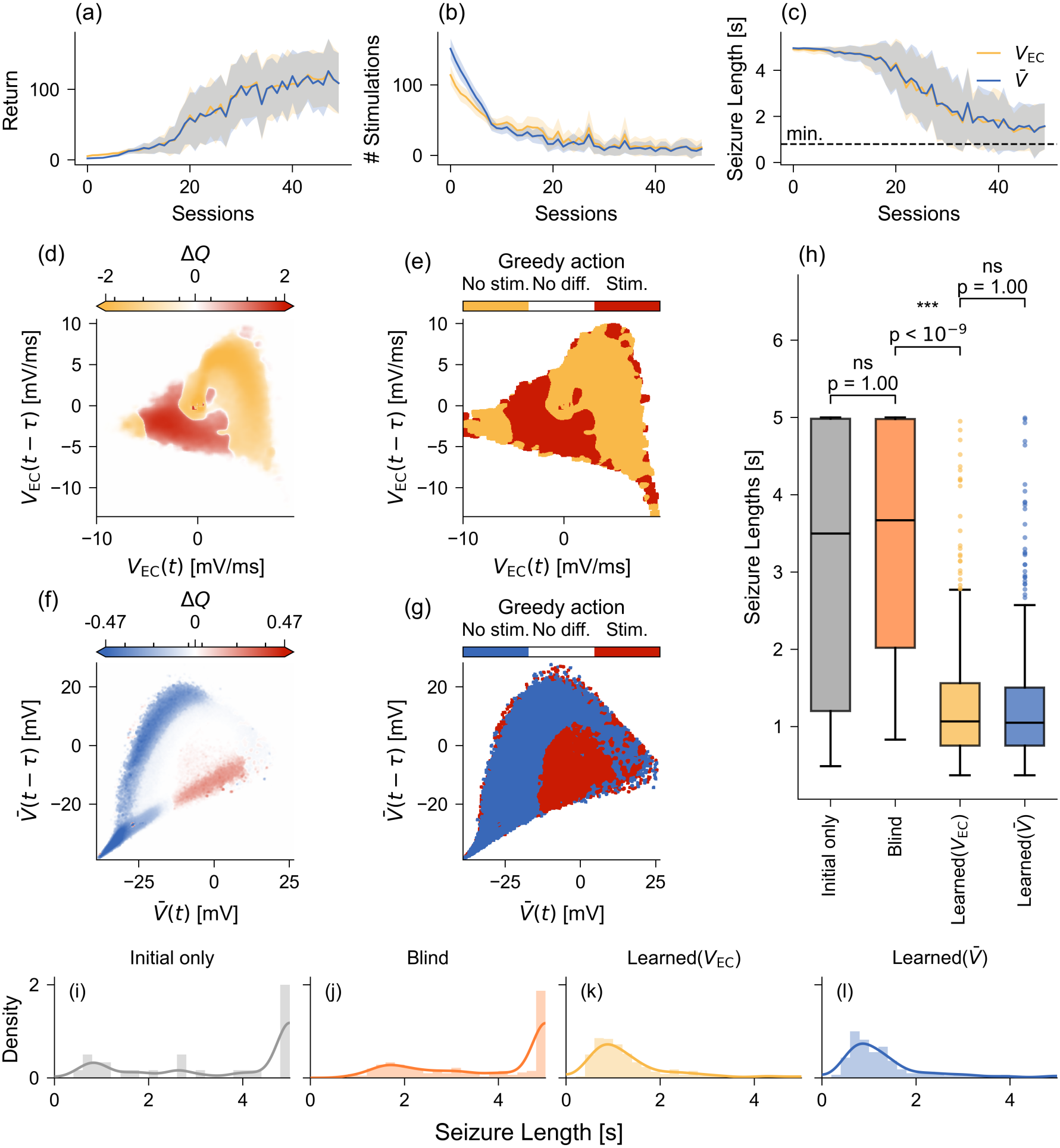
Learning and performance of the Q-learning agent on the two state variables. (a–c) Learning curves for the return, the number of stimuli and the seizure length, shown as mean ± SD. The dashed line is the estimated average minimum seizure duration (0.8 s), which equals the average activation delay. Yellow represents the agent trained on the extracellular potential proxy (*V*_EC_), blue the agent trained on the membrane potential 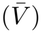. (d, f) Action-value differences Δ*Q* = *Q*_stim_ − *Q*_no stim_ at every state, using *V*_EC_ (d) or 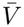 (f). The scale is symmetric-logarithmic and clipped at the 99th percentile of the non-zero values. (e, g) The greedy policies, i.e. the sign of Δ*Q*. (h) Comparison of seizure durations across four conditions: initial trigger only, blind (state-independent) stimulation, and policies learned using *V*_EC_ and 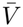 as state variables. Medians (horizontal lines), interquartile ranges (boxes), whiskers extending to ±1.5 IQR, and outliers (dots) are shown. (i–l) Histograms of the same four distributions, together with kernel density estimates.

During training, the agent determined the action-values of each state. When training was finished, these values were fixed and the agent used greedy action selection: the action with the higher value was chosen. Figure 4(d–g) illustrates the state spaces constructed using either 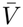 or *V*_EC_ as the state variable, coloured according to the learned action-value preferences. States in which stimulation is preferred are highlighted in red, while states with equal action-values are shown in white. States in which the no-stimulation action (*A*_0_) is preferred retain the colour associated with the corresponding state variable (yellow for *V*_EC_ and blue for 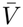). In Fig. 4 (d) and (f), the magnitude of the difference between the action-values is indicated by the strength of the shading, the darker the colour of a state, the stronger the agent’s preference, while lighter shades indicate uncertainty. Fig. 4 (e) and (g) show the corresponding binary policies, which indicate the action selected in each state during testing.

The shape of the red region in Fig. 4 (d–e) is complex and does not simply occupy a segment of the attractor. This indicates that the phase of *V*_EC_ alone does not provide sufficient information to determine an optimal stimulation policy. In this experiment, the optimal stimulation phase and amplitude are interdependent and jointly determine the outcome of stimulation.

We generated 30 new seizures for testing and ran the agent 10 times on each. We measured its performance by the duration of the seizures and compared the results with two control conditions. In the first control, the network evolved after the initial trigger without intervention. In the second, we used an agent with a state-independent policy that stimulated six times at random instants within the first 1.13 s, based on the results of the trained agents (see Table C7). The seizure durations under these conditions are shown in Fig. 4 (h–l).

Seizure length did not differ significantly between the networks receiving the initial trigger only and those under the blind agent (median over networks 3.5 s vs 3.8 s; two-sided Mann–Whitney U on one median per network, *n* = 30 per group, Holm-corrected over all six pairs, *p* = 1.0). Note the high number of seizures that lasts until the end of the simulations in both cases (Fig. 4) (i,j). Both learned policies shortened seizures to about a third of either control (median 1.11 s for *V*_EC_, 1.13 s for 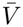; *p <* 10^−9^ against the blind agent and *p* = 0.001 against the trigger-only condition for both). The two learned policies did not differ significantly from each other (*p* = 1.0), suggesting that the noisier, but experimentally available *V*_EC_ carries enough information for the agent to control the seizure as well as the membrane potential does.

Fig. 5 demonstrates how a single, well-timed stimulus can terminate a seizure. It shows the activity of the same network in the time domain and in the phase plane following two stimulations delivered at different times, with the unstimulated network trajectory shown in the background for reference. In this example, the poorly timed stimulation prolonged the seizure until the end of the simulation, whereas the well-timed stimulation terminated it within a single oscillation cycle. Notably, the successful stimulation (red arrow in panel (f)) displaced the trajectory out-ward from the periodic attractor compared with the unstimulated trajectory over the same interval (yellow arrow). In contrast, the unsuccessful stimulation (orange arrow in panel (c)) displaced the trajectory inward, toward the periodic attractor, compared with the corresponding unstimulated trajectory (yellow arrow).

**Fig 5:**
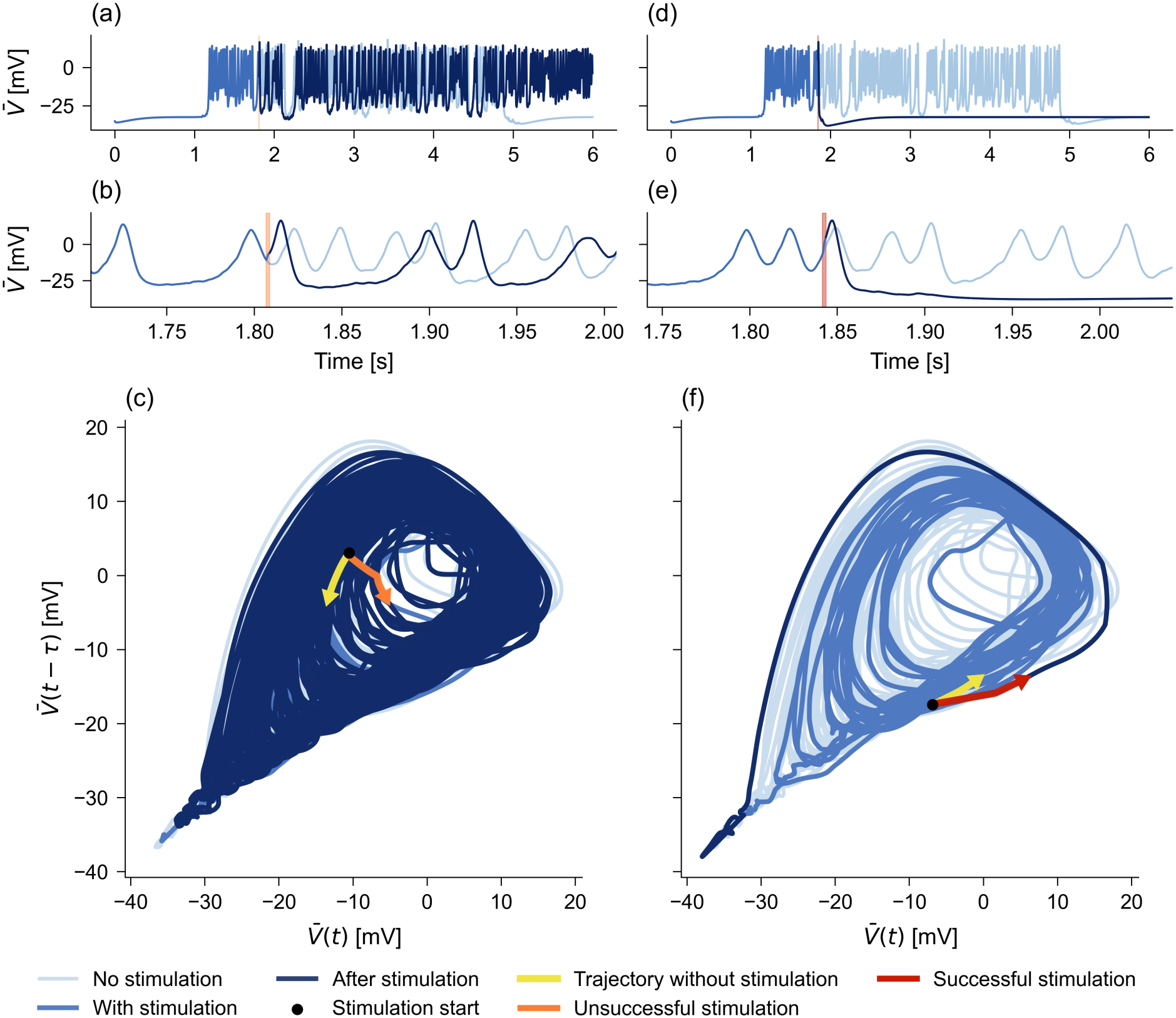
Comparison of the effects of unsuccessful and successful stimulation on network dynamics. The left column (a–c) shows an unsuccessful stimulation, which failed to terminate and even prolonged the seizure, while the right column (d–f) shows a successful stimulation delivered by the trained Q-learning agent, which terminated the seizure within a single oscillation cycle. The original seizure trajectory following the initial stimulus is shown in light blue in the background in both conditions. Panels (a,d) show the effects of stimulation in the time domain, with (b,e) providing magnified views around the stimulation time. Panels (c,f) show the effects of stimulation in the delay-embedded state space. Note that the well-timed stimulation (f) moved the system state outward from the periodic attractor, whereas the unsuccessful stimulation (c) moved it inward.

## 4 Discussion

Our study demonstrated how a state-dependent stimulation protocol can be learned by a reinforcement learning agent and used to terminate seizure-like activity efficiently in Morris–Lecar networks.

Jirsa et al. (2014) proposed that ictal and normal brain states have to coexist for seizures to occur. A mathematical model capturing this phenomenon has to exhibit a bistable regime of rest and oscillation. Additionally, they argued that the invariant properties of seizures are likely to arise from conserved bifurcations at seizure onset and offset.

These two dynamical constraints, bistability and typical bifurcations, were present in our Morris–Lecar model. The model showed bistability for 41.4*μ*A*/*cm^2^ ≤ *I* ≤ 44.34*μ*A*/*cm^2^, where the equilibrium branch could represent the normal state and the oscillatory trajectory the ictal state. In the second bistable region, 45.17 ≤ *I* ≤ 47.75*μ*A*/*cm^2^, the equilibrium branch could correspond to a depolarization block and should therefore be avoided. A possible mechanism of seizure generation in our system is that one neuron enters the basin of attraction of the limit cycle due to the initial stimulus and then spreads the activity to neighbouring neurons through the synaptic alpha functions. This could result in synchronous, seizure-like network activity. Spontaneous seizure termination may occur when the input to the neurons decreases sufficiently, so that the oscillatory attractor disappears through a homoclinic bifurcation and the cells return to the resting state. It is also possible that cells cross the separatrix in the bistable regime as their state variables change, and thus enter the basin of attraction of the stable fixed point. As more cells return to rest, the probability of neurons exciting each other decreases and the system can potentially recover.

Beyond its dynamical properties, the model produces electrophysiological signals that qualitatively resemble those observed during epileptic seizures. It exhibits strongly synchronized (Golomb–Rinzel synchrony index of 0.730) yet irregular spikes and high-frequency discharges characteristic of seizure onset. Trajectories in the phase plane also have a structure similar to that seen in Jirsa et al. (2014). This is especially important, as the input to our agent is a point in that space.

Bistability may not only explain why the same network shows different behaviours, but it is also a property that can be exploited when establishing a stimulation strategy. Rinzel (1985) showed that the Morris–Lecar neuron has bistable regimes, and that in these regimes a properly timed stimulation pulse can either induce persistent firing or redirect the system to the resting state. Jirsa et al. (2014) suggested that a separatrix has to be crossed for seizure termination.

Although our model has a higher-dimensional state space, its dynamics resemble those of the LCs class described by Saggio et al. (2020) and Szuromi et al. (2023). The basin of attraction of the fixed point wraps around the limit cycle, so stimuli that displace trajectories outward can drive the system into the basin of attraction of the fixed point and thereby terminate the seizure. Consistent with the findings of Szuromi et al. (2023), low-amplitude stimulation is effective only in specific regions of the state space. Here, we demonstrate that reinforcement learning can acquire this information through trial and error, learning to deliver stimulation in the appropriate states of the system.

Classical control-theoretic approaches have also been applied to suppress seizures (Kramer et al. 2006). A system of 14 coupled differential equations was introduced to model cortical epileptic activity, and three different feedback controller algorithms were applied to suppress seizures. Note that such control requires the delivery of either a negative or a positive stimulus at every time instant. Moreover, the controller suppressed the seizure only during the period when it was switched on, unlike our approach, which was able to terminate the seizure completely.

Suffczynski et al. (2004) and Taylor et al. (2014) examined the state dependence of the response of an epileptic system to stimuli in computational models of absence epilepsy. Taylor et al. concluded that successful stimulation depends on the system’s position in state space, where the basin of attraction of the resting state forms a complex, four-dimensional structure. This result highlights the need for an algorithm that can navigate such a complex state-space structure and learn state-dependent stimulation policies with-out knowing a priory the underlying dynamical structure.

Nevertheless, the study has some limitations. As much as the computational model helps the training of the algorithm, it is a strong simplification of a real system. It is a useful tool for gaining insight into the learning process, but the ultimate goal would be to train the agent on real data. The large number of training runs and episodes is also a constraint that we have to overcome if we wish to apply the method to patients with fewer epileptic episodes. Although the algorithm successfully identified a state-dependent protocol, the parameters of the stimulation waveform were fixed. In the future, the algorithm could learn these parameters as well and generate different waveforms for different states. Overall, this study can be a step towards exploiting the dynamical features of seizures combined with learning algorithms in a personalized, state-dependent stimulation protocol.

## 5 Conclusion

We built a network of Morris–Lecar neurons that was capable of generating seizure-like activity patterns, and we trained a semi-gradient Q-learning agent to stimulate the network in order to reduce seizure durations. The median seizure duration decreased from 3.5 s without intervention to 1.13 s and 1.11 s for agents trained using membrane potential and extracellular potential proxy signals, respectively. The seizure durations under a blindly stimulating control agent were indistinguishable from those of the original seizures after the initial stimulus. The input of the agent was a single scalar signal and one delayed copy of it, and an agent given a proxy for the extracellular potential performed no worse than one given the membrane potential directly. Reinforcement learning can therefore recover a state-dependent stimulation policy from the kind of minimal, realistically measurable input an implanted device would have.

## Declarations

## Acknowledgements

The authors are grateful to Professor Péter Érdi for invaluable insights during the preparation of this manuscript.

## Funding

This work was supported by the TECH-2024-020 grant of the HUN-REN Hungarian Research Network.

## Competing interests

The authors have no competing interests to declare.

## Ethics approval and consent to participate

Not applicable.

## Consent for publication

Not applicable.

## Data availability

Not applicable.

## Materials availability

Not applicable.

## Code availability

Upon request.

## Author contributions

Conceptualization: ZS and TK. Simulations and analysis: VH, MS, ZS. Founding acquisition: MS, TK, ZS. Manuscript drafting: VH, ZS, KT. All authors approved the manuscript.

## Appendix A Parameters

## Appendix B Linear stability of the Morris–Lecar equilibria

## Appendix C RL agent testing result

**Table A1:**
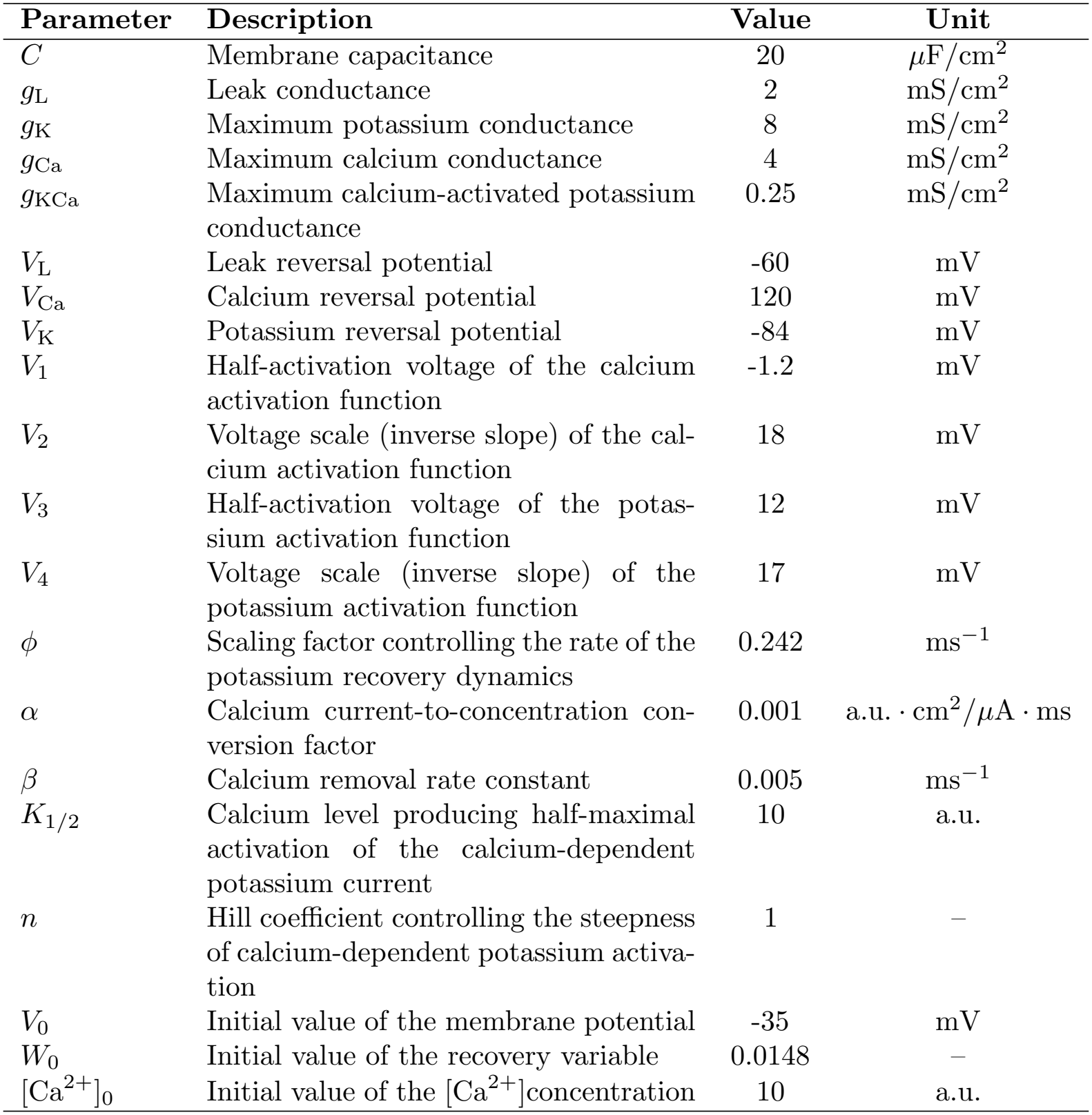
Parameters of the three-dimensional Morris–Lecar neuron model. All neurons in the network shared the same parameters. The unit of the calcium concentration [Ca^2+^] was arbitrary (a.u.), and the parameters *α, K*_1*/*2_ and [Ca^2+^]_0_ inherited this unit.

| Parameter | Description | Value | Unit |
|---|---|---|---|
| *C* | Membrane capacitance | 20 | µF/cm<sup>2</sup> |
| *g*<sub>L</sub> | Leak conductance | 2 | mS/cm<sup>2</sup> |
| *g*<sub>K</sub> | Maximum potassium conductance | 8 | mS/cm<sup>2</sup> |
| *g*<sub>Ca</sub> | Maximum calcium conductance | 4 | mS/cm<sup>2</sup> |
| *g*<sub>KCa</sub> | Maximum calcium-activated potassium conductance | 0.25 | mS/cm<sup>2</sup> |
| *V*<sub>L</sub> | Leak reversal potential | -60 | mV |
| *V*<sub>Ca</sub> | Calcium reversal potential | 120 | mV |
| *V*<sub>K</sub> | Potassium reversal potential | -84 | mV |
| *V*<sub>1</sub> | Half-activation voltage of the calcium activation function | -1.2 | mV |
| *V*<sub>2</sub> | Voltage scale (inverse slope) of the cal-cium activation function | 18 | mV |
| *V*<sub>3</sub> | Half-activation voltage of the potas-sium activation function | 12 | mV |
| *V*<sub>4</sub> | Voltage scale (inverse slope) of the potassium activation function | 17 | mV |
| *ϕ* | Scaling factor controlling the rate of the potassium recovery dynamics | 0.242 | ms<sup>−1</sup> |
| *α* | Calcium current-to-concentration con-version factor | 0.001 | a.u. · cm<sup>2</sup>/µA · ms |
| *β* | Calcium removal rate constant | 0.005 | ms<sup>−1</sup> |
| *K*<sub>1/2</sub> | Calcium level producing half-maximal activation of the calcium-dependent potassium current | 10 | a.u. |
| *n* | Hill coefficient controlling the steepness of calcium-dependent potassium activa-tion | 1 | – |
| *V*<sub>0</sub> | Initial value of the membrane potential | -35 | mV |
| *W*<sub>0</sub> | Initial value of the recovery variable | 0.0148 | – |
| [Ca<sup>2+</sup>]<sub>0</sub> | Initial value of the [Ca<sup>2+</sup>]concentration | 10 | a.u. |

**Table A2:**
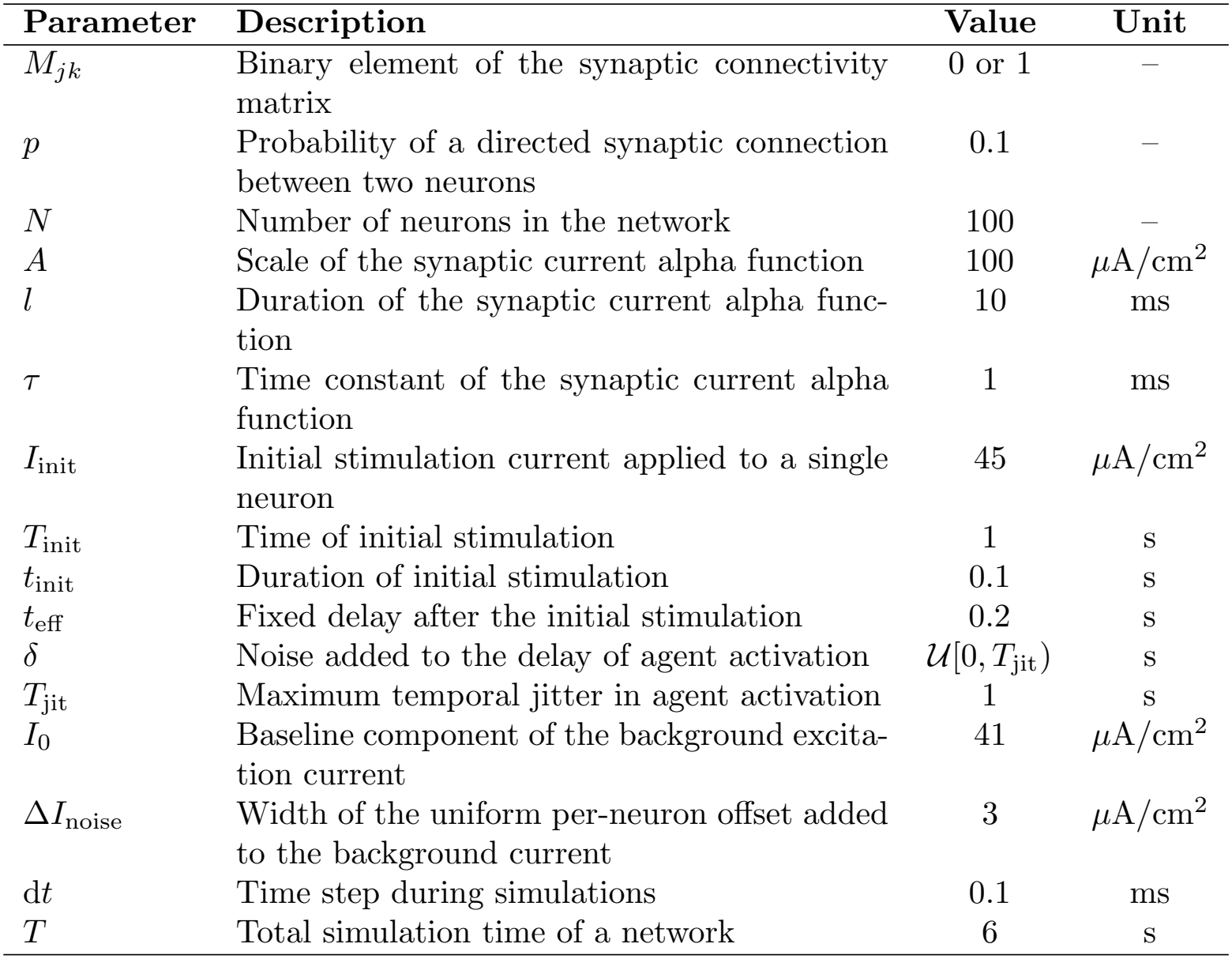
Parameters of the network and simulations.

| **Parameter** | **Description** | **Value** | **Unit** |
|---|---|---|---|
| *M*<sub>*jk*</sub> | Binary element of the synaptic connectivity matrix | 0 or 1 | – |
| *p* | Probability of a directed synaptic connection between two neurons | 0.1 | – |
| *N* | Number of neurons in the network | 100 | – |
| *A* | Scale of the synaptic current alpha function | 100 | *μ*A/cm<sup>2</sup> |
| *l* | Duration of the synaptic current alpha func-tion | 10 | ms |
| *τ* | Time constant of the synaptic current alpha function | 1 | ms |
| *I*<sub>init</sub> | Initial stimulation current applied to a single neuron | 45 | *μ*A/cm<sup>2</sup> |
| *T*<sub>init</sub> | Time of initial stimulation | 1 | s |
| *t*<sub>init</sub> | Duration of initial stimulation | 0.1 | s |
| *t*<sub>eff</sub> | Fixed delay after the initial stimulation | 0.2 | s |
| *δ* | Noise added to the delay of agent activation | 𝒰[0, *T*<sub>jit</sub>) | s |
| *T*<sub>jit</sub> | Maximum temporal jitter in agent activation | 1 | s |
| *I*<sub>0</sub> | Baseline component of the background excita-tion current | 41 | *μ*A/cm<sup>2</sup> |
| Δ*I*<sub>noise</sub> | Width of the uniform per-neuron offset added to the background current | 3 | *μ*A/cm<sup>2</sup> |
| d*t* | Time step during simulations | 0.1 | ms |
| *T* | Total simulation time of a network | 6 | s |

**Table A3:**
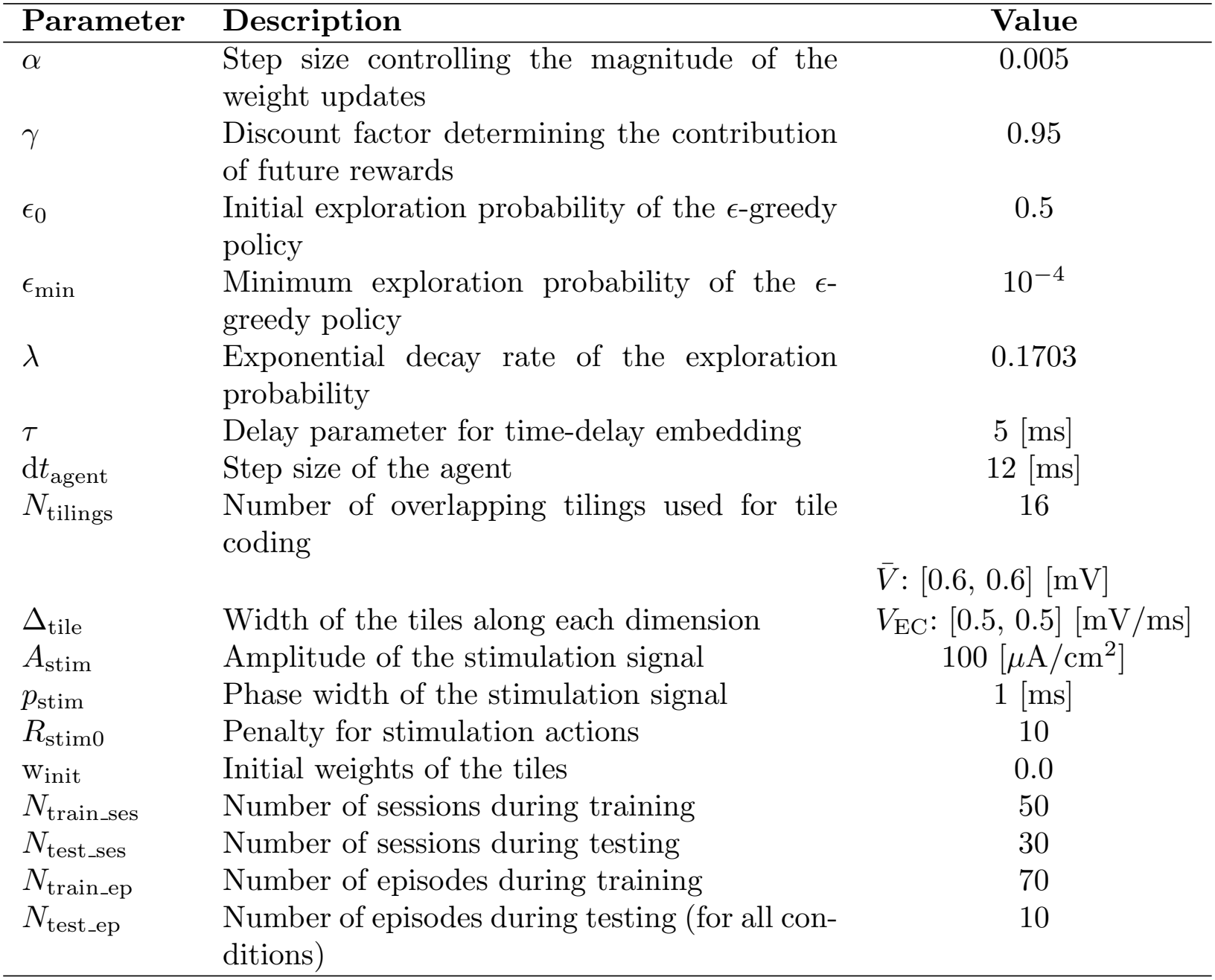
Parameters of the reinforcement learning algorithm.

| Parameter | Description | Value |
|---|---|---|
| α | Step size controlling the magnitude of the weight updates | 0.005 |
| γ | Discount factor determining the contribution of future rewards | 0.95 |
| ϵ<sub>0</sub> | Initial exploration probability of the ϵ-greedy policy | 0.5 |
| ϵ<sub>min</sub> | Minimum exploration probability of the ϵ-greedy policy | 10<sup>−4</sup> |
| λ | Exponential decay rate of the exploration probability | 0.1703 |
| τ | Delay parameter for time-delay embedding | 5 [ms] |
| dt<sub>agent</sub> | Step size of the agent | 12 [ms] |
| N<sub>tilings</sub> | Number of overlapping tilings used for tile coding | 16 |
| Δ<sub>tile</sub> | Width of the tiles along each dimension | V̄: [0.6, 0.6] [mV]<br>V<sub>EC</sub>: [0.5, 0.5] [mV/ms] |
| A<sub>stim</sub> | Amplitude of the stimulation signal | 100 [µA/cm<sup>2</sup>] |
| p<sub>stim</sub> | Phase width of the stimulation signal | 1 [ms] |
| R<sub>stim0</sub> | Penalty for stimulation actions | 10 |
| w<sub>init</sub> | Initial weights of the tiles | 0.0 |
| N<sub>train_ses</sub> | Number of sessions during training | 50 |
| N<sub>test_ses</sub> | Number of sessions during testing | 30 |
| N<sub>train_ep</sub> | Number of episodes during training | 70 |
| N<sub>test_ep</sub> | Number of episodes during testing (for all con-ditions) | 10 |

**Table B4:** Equilibrium points and corresponding eigenvalues of the three-dimensional Morris–Lecar model at *I* = *I*_0_.

| No. | *V*<sup>\*</sup> | *W*<sup>\*</sup> | [Ca<sup>2+</sup>]<sup>*</sup> | Eigenvalues (*λ*) |
|---|---|---|---|---|
| #1 | −21.562 | 0.019 | 10.677 | 0.091 |
|  |  |  |  | −0.326 |
|  |  |  |  | −0.004 |
| #2 | −34.078 | 0.004 | 3.113 | − 0.034 |
|  |  |  |  | − 0.489 |
|  |  |  |  | − 0.007 |
| #3 | 3.459 | 0.268 | 58.421 | 0.007 ± 0.369*i* |
|  |  |  |  | −0.005 |

**Table B5:** Eigenvalues and state variables at different input currents near the bifurcation point (*I*_b_ = 39.85 *μ*A*/*cm^2^). The eigenvalues indicate that the equilibrium remains a hyperbolic saddle as the input current approaches the bifurcation point.

| *I* [*μ*A/cm<sup>2</sup>] | *T* [s] | *f* [Hz] | *λ*<sub>1</sub> | *λ*<sub>2</sub> | *λ*<sub>3</sub> | *V* | *W* | Ca |
|---|---|---|---|---|---|---|---|---|
| 40.40 | 0.1829 | 5.4675 | 0.098369 | -0.320559 | -0.003990 | -21.105 | 0.019944 | 11.143 |
| 40.21 | 0.2505 | 3.9920 | 0.100553 | -0.318870 | -0.004009 | -20.960 | 0.020265 | 11.287 |
| 40.00 | 0.4181 | 2.3918 | 0.102870 | -0.317090 | -0.004029 | -20.820 | 0.020600 | 11.441 |
| 39.96 | 0.4739 | 2.1101 | 0.103243 | -0.316805 | -0.004032 | -20.797 | 0.020665 | 11.466 |
| 39.93 | 0.5427 | 1.8426 | 0.103574 | -0.316552 | -0.004034 | -20.776 | 0.020714 | 11.488 |
| 39.91 | 0.6288 | 1.5903 | 0.103857 | -0.316336 | -0.004037 | -20.758 | 0.020757 | 11.507 |
| 39.89 | 0.7387 | 1.3537 | 0.104086 | -0.316161 | -0.004038 | -20.744 | 0.020790 | 11.522 |
| 39.87 | 0.8819 | 1.1339 | 0.104257 | -0.316030 | -0.004040 | -20.733 | 0.020800 | 11.534 |
| 39.86 | 1.0749 | 0.9303 | 0.104371 | -0.315944 | -0.004041 | -20.760 | 0.020830 | 11.541 |
| 39.85 | 7.9961 | 0.1251 | 0.104457 | -0.315878 | -0.004041 | -20.721 | 0.020847 | 11.547 |

**Table B6:**
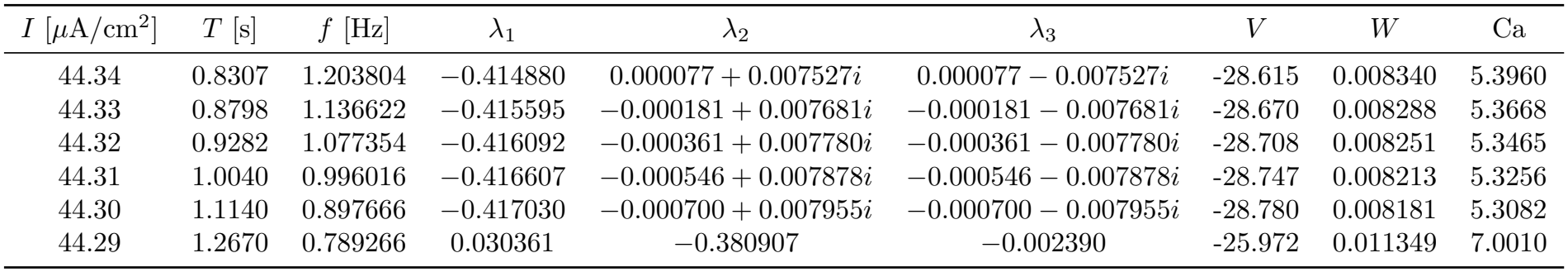
Eigenvalues and state variables at different input currents near the bifurcation point (*I*_b_ = 44.29 *μ*A*/*cm^2^). The eigenvalues indicate a transition from a stable focus to an unstable saddle point as the input current approaches the bifurcation point.

**Table C7:**
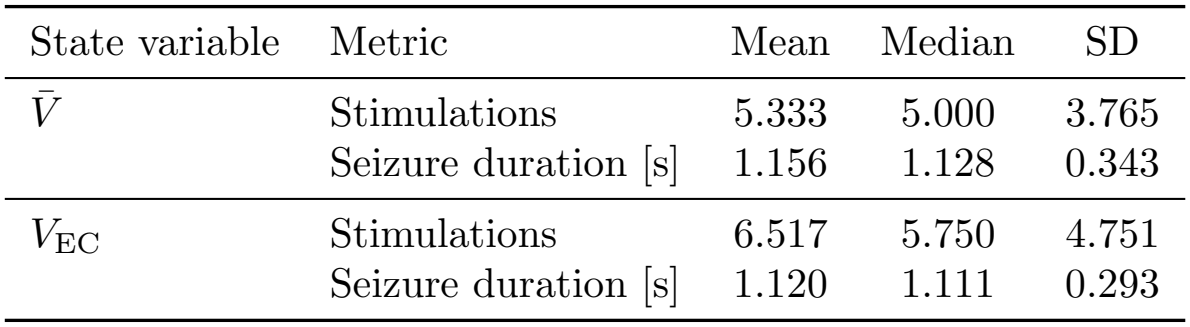
Descriptive statistics of the number of stimulations and seizure duration for agents trained using membrane potential 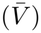 and extracellular potential (*V*_EC_). For each of the 30 sessions, the median was first calculated across the 10 episodes. The reported statistics were then computed from the resulting 30 group-level medians.

| State variable | Metric | Mean | Median | SD |
|---|---|---|---|---|
| *V̄* | Stimulations | 5.333 | 5.000 | 3.765 |
|  | Seizure duration [s] | 1.156 | 1.128 | 0.343 |
| *V*<sub>EC</sub> | Stimulations | 6.517 | 5.750 | 4.751 |
|  | Seizure duration [s] | 1.120 | 1.111 | 0.293 |

**Fig B1:**
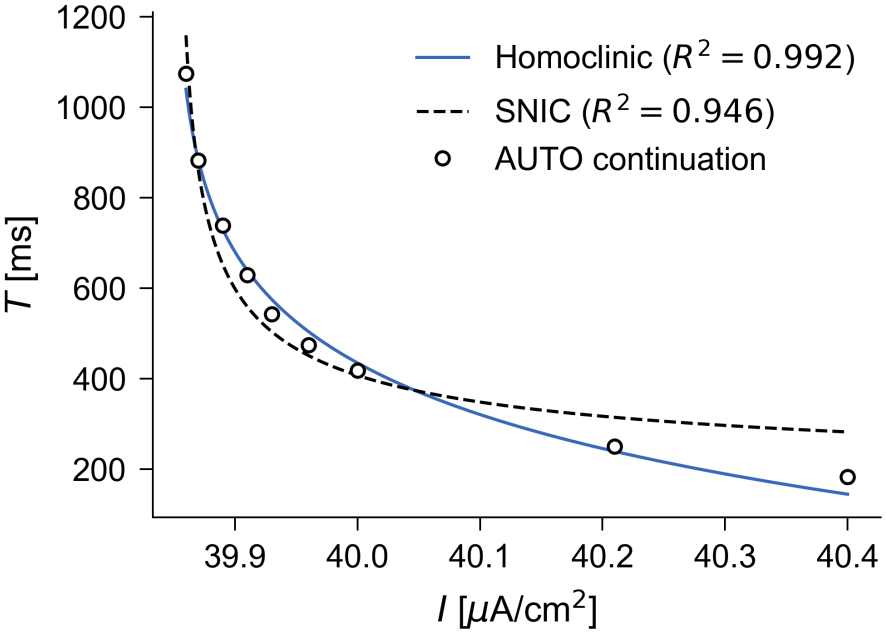
Fitting of the characteristic functions for a saddle-homoclinic bifurcation (HC) and for a saddle-node on invariant circle (SNIC) bifurcation. The larger *R*^2^ value indicates a better fit, which implies the type of bifurcation around *I* = 44.29*μ*A*/*cm^2^.

**Fig B2:**
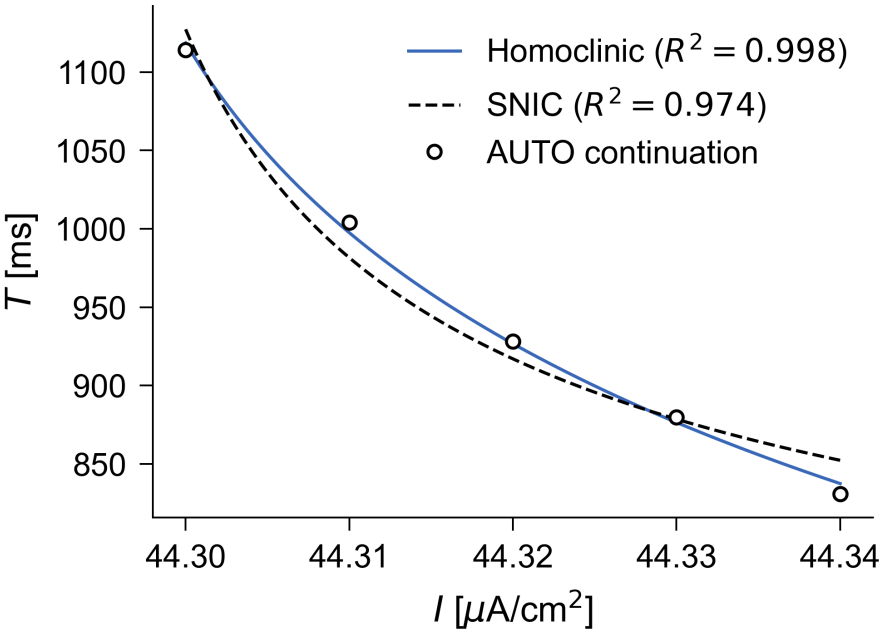
Fitting of the characteristic functions for a saddle-homoclinic bifurcation (HC) and for a saddle-node on invariant circle (SNIC) bifurcation. The larger *R*^2^ value indicates a better fit, which implies the type of bifurcation around *I* = 39.85*μ*A*/*cm^2^.

## Notes

### Competing Interest Statement

The authors have declared no competing interest.

